# Decomposing the effects of α-tACS on brain oscillations and aperiodic 1/f activity

**DOI:** 10.1101/2023.10.25.563756

**Authors:** Florian H. Kasten, René Lattmann, Daniel Strüber, Christoph S. Herrmann

## Abstract

**Background:** Aftereffects of transcranial alternating current stimulation (tACS) are usually analyzed with a focus on the individual frequency band, thereby neglecting broadband spectral components. Recently, it was shown that the broadband spectrum, which exhibits a 1/f-like characteristic, is functionally relevant.

**Objective/Hypothesis:** The goal of this study was a spectral parameterization of brain activity into oscillatory alpha activity and aperiodic 1/f components before and after tACS and sham stimulation. It was expected that the broadband spectrum will not be differentially influenced by 20-min of tACS at individual α-frequency (IAF) in comparison to sham. Additionally, it was expected that the tACS aftereffect on the α-band can still be observed, even when controlling for 1/f activity differences.

**Methods:** We performed a re-analysis of a recently published resting-state tACS-magnetoencephalography (MEG) data set. Parameterization of the frequency spectrum was computed with the fitting-oscillations-and-one-over-F (FOOOF) algorithm. The intercept as well as the slope parameter of the aperiodic 1/f fit was extracted. Data was analyzed in sensor space with a focus on magnetometers. Comparison of changes in α-band power and 1/f activity was performed with non-parametric cluster-based random permutation tests.

**Results:** The tACS aftereffect survived the 1/f-correction. The previously observed natural rise in alpha oscillations over time independent of experimental conditions could not be replicated. However, differences in aperiodic parameters over time were observed. Especially, the intercept parameter increased from pre to post stimulation to a similar degree in both conditions.

**Conclusion:** It is imperative to correct for the aperiodic 1/f spectral component when analyzing aftereffects of brain stimulation on brain oscillations.

## 1 Introduction

Transcranial alternating current stimulation (tACS) has become a popular tool in neuroscience to non-invasively interfere with oscillations in the human brain [1,2]. This brain stimulation technique works via the application of sinusoidal currents between at least two electrodes placed on the scalp and is based on the core assumption that it modulates brain oscillations and their associated cognitive processes in a frequency-specific manner [1,3,4]. Interventional studies using tACS are particularly interesting as they allow to establish causal relationships between oscillations within particular frequency bands and cognitive functions [5].

Brain oscillations can be described as short-term periodic signals with specific frequencies which are typically divided into different frequency bands known as delta (0-4 Hz), theta (4-8 Hz), alpha (8-12 Hz), beta (12-30 Hz) and gamma (30-80 Hz) oscillations [5–7]. Those frequency bands usually contain oscillations with different amplitudes. That is, frequencies within the lower part of the spectrum have higher amplitudes while higher frequencies have lower amplitudes. This amplitude distribution across the frequency spectrum has a certain shape. It follows a so-called 1/f-like distribution. The term 1/f-*like* stems from the fact that the distribution does not usually follow a true 1/f function [8]. Interestingly, also the non-oscillatory brain activity follows a 1/f-distribution and has been referred to as scale-free [9,10] or fractal (i.e. statistically self-similar [8]) brain activity. This stems from the fact that underlying processes are proposed to be scale-invariant in that they are devoid of periodicity [9]. This phenomenon is ubiquitous in nature and can be found in many different systems [11]. In the following, we will use the term *1/f brain activity* for the sake of simplicity. This aperiodic component of brain activity can be represented as a line when depicted in a double logarithmic plot and is characterized by the line’s parameters *slope* (synonymous with the term ‘exponent’) and *intercept* (synonymous with the term ‘offset’), respectively. While previous studies regarded 1/f brain activity as pure noise, recent advances have put forward the notion that 1/f brain activity is in fact functionally relevant [9,10,12–15]. For example, by simulating local field potentials (LFPs), Gao et al. [12] could show that changes in the excitation-inhibition ratio result in changes of the aperiodic 1/f brain activity. The excitation-inhibition ratio of synaptic inputs is an essential requirement for neural homeostasis [16]. Building on the modeling and invasive animal work of Gao et al. [12], Waschke et al. [15] demonstrated in humans that the spectral slopes of non-invasive EEG recordings reflect not only general changes in the balance of excitatory to inhibitory neural activity but also behaviorally relevant variations of the attentional focus in a target-detection task. Additionally, Voytek et al. [14] analyzed power spectra from electrocorticograms (ECoG) of participants of different age groups. They could show that the slope parameter of the aperiodic 1/f brain activity is lower for older age in comparison to younger age. This was proposed to be a result of increasing noise levels in cognitive processing in older age [14].

Recently, Ouyang et al. [13] studied the relationship between resting-state EEG activity and the efficiency of cognitive functioning. The authors segregated aperiodic 1/f brain activity from alpha oscillations to explain variance of cognitive processing speed using structural equation modelling. They found that variance in aperiodic 1/f brain activity, but not alpha oscillations, explained variance in cognitive processing speed. Thus, once aperiodic 1/f brain activity is taken into account, it can explain significant amounts of variance which otherwise would have been mistaken as originating from differences in oscillatory amplitudes.

Given the evidence for a functional role of aperiodic 1/f brain activity in cognitive processing and the resulting necessity of isolating this component from oscillatory activities, the question arises whether tACS only modulates oscillatory amplitudes or whether it may modulate aperiodic 1/f brain activity as well. Further, the question arises whether established tACS aftereffects in the alpha band [17– 21] are still observable once aperiodic 1/f brain activity is accounted for. To address these issues in the present study, we used an existing dataset of tACS aftereffects on alpha oscillations and dissociated tACS effects on oscillations from those on aperiodic 1/f brain activity by separately parameterizing the different components of the frequency spectrum. Our results show that tACS aftereffects on brain oscillations are still observed after removing aperiodic 1/f brain activity. Nevertheless, there are changes in the aperiodic 1/f parameters slope and intercept across time. Disentangling these effects may provide further insights into the exact mechanisms underlying tACS aftereffects.

## 2 Materials and methods

### 2.1 Participants

The analyses in the current study were carried out on a pre-existing tACS-MEG dataset [22]. TACS at individual alpha frequency (IAF) or sham stimulation were administered during two experimental sessions on separate days. Participants (N = 19, 25 ± 3 years, 11 females), received both stimulation conditions in random order. They were without history of neurological or psychiatric disease and medication-free at the day of the experiment. They were right-handed according to the Edinburgh Handedness-Inventory [23], non-smokers and had normal or corrected-to-normal vision. Written informed consent was obtained from all subjects, both experiments were approved by the Commission for Research Impact assessment and Ethics at the University of Oldenburg and complied with all relevant ethical regulations.

### 2.2 Magnetoencephalogram (MEG)

MEG signals were sampled at 1 kHz using a 306-channel whole-head MEG system (Elekta Neuromag Triux System, Elekta Oy, Helsinki, Finland), inside a magnetic shielding room (MSR; Vacuumschmelze, Hanau, Germany). Participants’ head-position was monitored using five head-position indicator (HPI) coils. Coil positions and participants’ head shapes were digitized using a Polhemus Fastrak (Polhemus, Colchester, VT, USA).

### 2.3 Electrical Stimulation

TACS was administered using rectangular, surface-conductive rubber electrodes over locations Cz (7 x 5 cm) and Oz (4 x 4 cm) of the international 10-10 system. They were attached with an electrically conductive, adhesive paste (ten20 paste, Weaver & Co, Aurora, CO, USA). The stimulation waveform was digitally generated in Matlab (MATLAB 2016a, The Math Works Inc., Natick, MA, USA) and fed to the remote input of a constant current stimulator (DC Stimulator Plus, Neuroconn, Illmenau, Germany) via a digital-to-analog converter (NI-USB 6251, National Instruments, Austin, TX, USA). Stimulation currents were guided into the MSR via the MRI extension-kit of the stimulator. Impedances were kept below 20 kΩ, including two 5 kΩ resistors inside the stimulation cables. Participants received either 20 min of active tACS or sham stimulation (30-sec of tACS at the beginning of the stimulation period) at their individual α-frequency IAF, determined from a 3 min resting MEG prior to the main experiment. 10-min of MEG was recorded directly before and after stimulation. Participants were instructed to keep their eyes open during the recording and to perform a visual change detection task. A white fixation-cross on gray background was rear-projected onto a screen (distance: ∼100 cm) inside the MSR. Participants manually responded to a 500 ms rotation of the cross by 45°. Targets occurred with an SOA of 10-110-sec. Additional details on the experimental procedures, including debriefing and the assessment of adverse effects can be found in a previous publication based on the data [22].

## 2.4 Data analysis

### 2.4.1 Preprocessing

Spatiotemporal signal space separation [24,25] with standard settings (L_in_ = 8, L_out_ = 3, correlation limit = 0.98) was performed to suppress interference from external noise sources and to compensate for participants’ head movements.

Subsequent MEG preprocessing was carried out using Python 3.9 using MNE-Python 1.0 [26]. To facilitate computations, the analysis was focused on magnetometer channels only. MEG signals acquired during tACS application were discarded from the analysis due to the massive electromagnetic artifact contaminating the data [27]. Data was resampled to 256 Hz and filtered between 1 Hz and 40 Hz using a finite impulse response filter. Subsequently, independent component analysis was performed using the extended infomax algorithm. Components reflecting cardiac, eye-movement or -blink artifacts were visually identified and removed before back-projecting signals into sensor space. The data was segmented into non-overlapping 2-sec epochs. Artifacts remaining after ICA denoising were removed by a threshold-based artifact rejection discarding epochs with an amplitude exceeding 3000 fT. Power spectral density (4-sec zero padding, Hanning window) was computed on the first 260 artifact free segments in each of the two blocks.

### 2.4.2 Fitting oscillations and 1/f

To decompose power spectra into oscillatory and non-oscillatory 1/f brain activity we utilized the FOOOF algorithm as described by Donoghue and colleagues [28]. The algorithm is implemented in a Python toolbox with the same name (https://fooof-tools.github.io/fooof/).

The fooof algorithm was fitted over a frequency range of 1 Hz – 30 Hz with an unconstrained maximum number of gaussian peaks. No knee parameter was applied to the fit. The algorithm was fit for the average spectrum per channel and block of each participant and the intercept as well as the slope parameter of the aperiodic 1/f fit were extracted from the data. The aperiodic 1/f component of the spectrum was then subtracted from the full reconstructed spectrum in linear space to obtain a 1/f corrected spectrum.

### 2.4.3 Statistical analysis

Statistical analyses were performed in MNE-Python. Participants’ power in the individual α-frequency band (IAF ± 2 Hz) was extracted from the uncorrected and 1/f corrected spectra in each block and condition (pre-/post-tACS vs. sham). In addition, we extracted the individual slope and intercept of the aperiodic fits. All data were submitted to dependent samples permutation cluster t-tests [29] with 10,000 permutations. Specifically, we tested for changes in the outcome measures above from pre-to post-intervention separately for the tACS and sham conditions. In case these two tests both yielded significant results in the same direction, this would indicate a main effect of factor time with factor levels pre to post.

A main effect of stimulation (factor levels tACS and sham) was not assessed as it is by itself not indicative of an aftereffect of tACS. Such an aftereffect of tACS would require an interaction of the factors time and stimulation. This interaction was tested by assessing the difference between the pre-to post-differences of the conditions tACS and sham. All cluster p-values were additionally corrected for the three multiple comparisons (baseline vs. tACS, baseline vs. sham, tACS vs. sham) using Bonferroni correction.

## 3 Results

### 3.1 Aftereffect of tACS in the uncorrected spectrum

Consistent with previous source-level analyses of the data [22,30], we observed an increase in power in the α-frequency band (p_cluster_ < .001) from pre-to post-stimulation in the tACS condition (Fig. 1 a,b), that was absent in the sham condition (p_cluster_ = .25, Fig. 1 c,d) on the sensor-level. When directly comparing α-power change between tACS and sham, permutation cluster analysis revealed a significantly larger power increase in the α-band after tACS as compared to sham (p_cluster_ = .007, Fig. 1 e,f). This reflects the expected interaction of factors time and stimulation.

**Figure 1:**
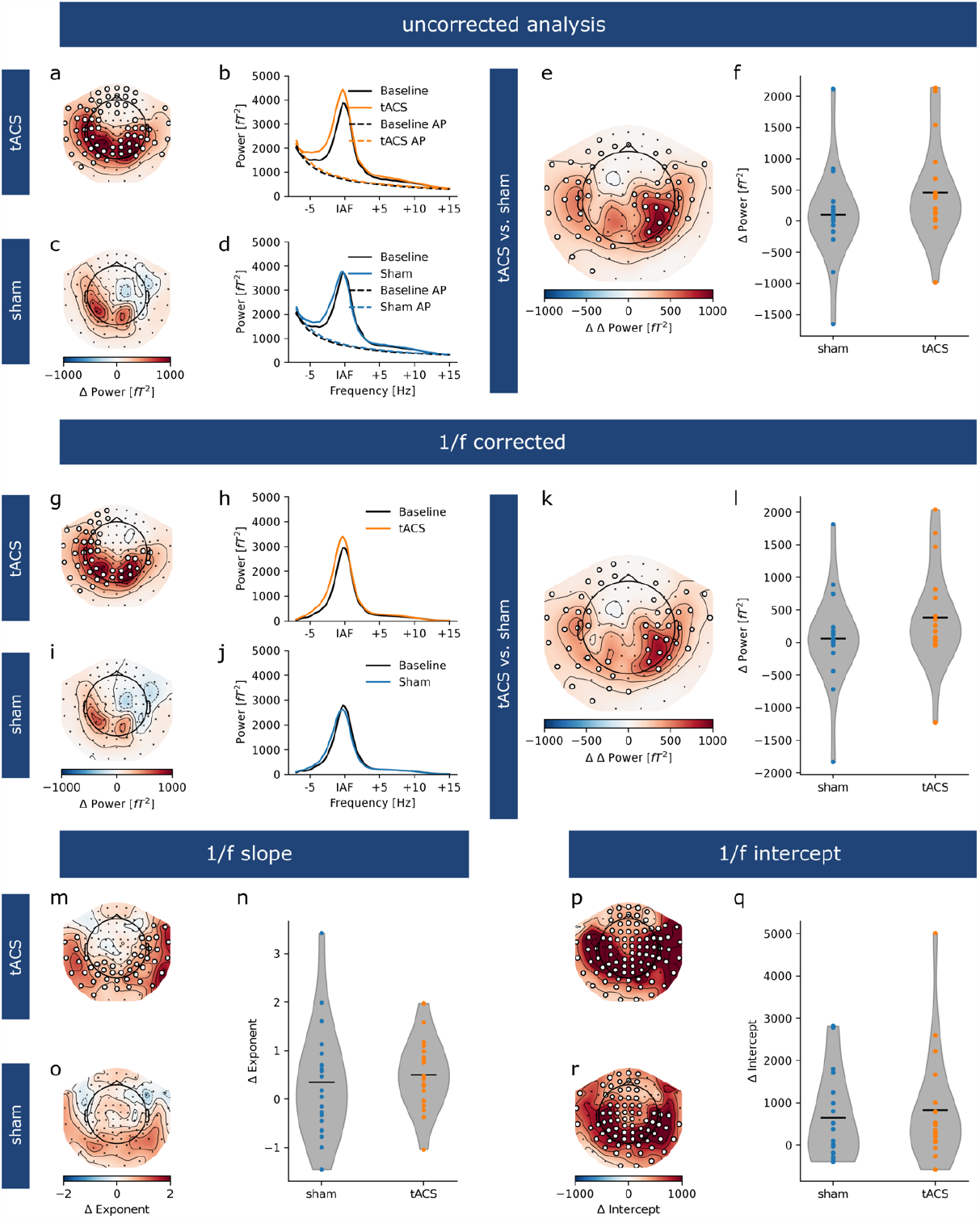
tACS modulates power in the α-band but not 1/f parameters. **a)** Topography of difference in spectral power in the α-band from pre-to post-stimulation in the tACS group. White markers indicate sensors with significant power increase. **b)** Average uncorrected power-spectra overlaid with the 1/f aperiodic fit (AP) before and after tACS. **c, d)** Same as in a, b for sham condition. **e)** Topography of difference of the difference in α-power from pre-to post-stimulation between tACS and sham [(tACS - baseline) - (sham - baseline)]. **f)** Violin plots depict distribution of pre-/post-stimulation α-power difference across participants. Black bars represent sample average. **g-l)** same as a-f after removing 1/f components from individual spectra. **m)** Topography of difference in aperiodic slope from pre- to post-tACS. **n)** Distribution of the change in aperiodic slope between tACS and sham conditions. **o)** same as m for sham condition. **p)** Topography of change in intercept of the aperiodic fit from pre-to post- tACS. **q)** Distribution of the change in aperiodic intercept between tACS and sham conditions. **r)** Same as p for sham condition.

### 3.2 Removing 1/f brain activity does not change the effect of tACS on α-oscillations

After removing 1/f brain activity from individual spectra, we observed nearly the same results as seen in the uncorrected data (Fig. 1 g-l). α-power increased significantly stronger after tACS as compared to sham (p_cluster_ < .011), reflecting the expected interaction of factors time and stimulation. While there was a significant increase in α-power from pre-to post-stimulation in the tACS group (p_cluster_ = .015), this effect was absent in the sham group (p_cluster_ = .56).

### 3.3 1/f slope and intercept parameters change over time, but seem unaffected by tACS

After decomposing power spectra into periodic and aperiodic components, we compared the slope and intercept parameters of the aperiodic fits between stimulation conditions. We observed a significant increase of the 1/f slope parameter from pre-to post in the tACS condition (p_cluster_ = .003, Fig. 1m), but not in the sham condition (p_cluster_ > .23, Fig. 1o). However, a direct comparison of these differences between tACS and sham (i.e, the interaction effect) did not differ significantly (p_cluster_ = 1, Fig. 1n). There is thus no clear evidence for an effect of tACS on the aperiodic slope, nor a consistent time-on-task effect.

In contrast, for the intercept of the aperiodic fit, we observed a significant increase from pre-to post-stimulation in both the tACS (p_cluster_ < .001, Fig 1p) and the sham group (p_cluster_ < .001, Fig 1r). The change of the intercepts did not differ between conditions (p_cluster_ = 1, Fig 1q), which reflects a main effect of the factor time (in absence of a main effect of the factor stimulation).

## 4 Discussion

### 4.1 Aftereffects of tACS in the uncorrected spectrum

We were able to replicate previous source-level analyses on the sensor-level MEG data, i.e. α -tACS upregulated MEG alpha oscillations which was not the case for sham stimulation.

### 4.2 Removing 1/f brain activity does not change the effect of tACS on α-oscillations

Estimating aftereffects of tACS on alpha power is typically done with a focus on the narrow frequency band around the IAF [9,17,19,20,22,28]. However, recent studies have provided first evidence that 1/f brain activity is functionally relevant for cognitive processing [9,10,12–15]. Thus, the important question arises whether tACS actually modulates only brain oscillations or whether it also affects 1/f brain activity.

The results of the present study demonstrate for the first time that aftereffects of tACS on alpha amplitude in comparison to sham stimulation are in fact present, even when correcting for 1/f brain activity. To date, no study explicitly decomposed periodic (i.e., oscillatory) and aperiodic 1/f components of the spectrum to analyze effects of tACS for each component separately. Although some studies controlled for 1/f brain activity on a general level by weighting higher frequencies more compared to lower frequencies [31,32], such frequency-weighting does not account for differences in aperiodic 1/f brain activity across participants. A different approach to control for 1/f brain activity is to detrend the signal by applying filters around the frequency band under study [33]. However, this procedure does not remove proportions of 1/f brain activity within the to-be-analyzed frequency band. As a result, spectral parameterization of individual frequency spectra, thereby accounting for interindividual differences in 1/f brain activity, may provide a valid alternative for estimating tACS aftereffects in alpha oscillations.

### 4.3 1/f slope and intercept parameters change over time, but seem unaffected by tACS

With regard to the slope parameter, the present study revealed that the change from pre-to post-stimulation is not significantly different between tACS and sham stimulation, that is, lacking an interaction of the factors time and stimulation. Therefore, the slope parameter seems not to be relevant for explaining the tACS aftereffect.

An interesting novel finding of the present study is the significant increase from baseline to post-stimulation in the intercept of aperiodic brain activity for both tACS and sham stimulation which did not differ between the stimulation conditions. Changes in aperiodic intercept have been related to changes in overall spiking-activity [14,34,35]. Thus, this result may indicate that tACS and sham stimulation affected the overall spiking activity by an increase in the aperiodic intercept to a similar degree. So far, research has shown that cortical excitability can increase naturally with prolonged phases of wakefulness [36]. However, Huber et al. [36] analyzed differences across multiple days while investigating sleep deprivation. Future research may examine whether natural cortical excitability changes are also present within shorter time periods.

Combining our results of a stimulation-unspecific increase in the aperiodic intercept over time and the 1/f-corrected power increase in the alpha band may help to further elucidate the mechanisms underlying tACS aftereffects. It is an often reported finding in studies on tACS aftereffects that alpha power not only increased after tACS but also after sham stimulation so that the overall tACS effect declined [17,19,37,38]. This natural rise of alpha power during sham stimulation has been attributed to conditions known to increase alpha power over time, such as vigilance decrement, time on task and mental fatigue [17,37], eyes closed vs. eyes open [19], or the level of ambient light [37,38]. Whereas these interpretations focus on the change in alpha power as a purely periodic phenomenon, the results of the present study point to another possible cause of the time-on-task-effect, reflected by an increase of the aperiodic 1/f intercept. An increase from pre to post intercept could be mistaken for an increase in oscillatory alpha power over time if only the pre to post peak values of the 1/f-uncorrected data are considered. Fig. 2a shows a time-on-task-effect for the sham condition as a function of an increase in the 1/f intercept parameter, that is, without any actual changes in brain oscillations. In contrast, the tACS pre to post effect reflects a combined increase in the 1/f intercept parameter and oscillatory power that can only be deciphered from 1/f-corrected data (Fig. 2b).

**Figure 2:**
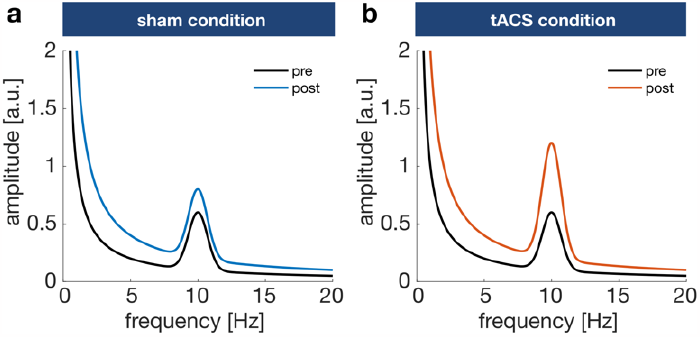
Schematic representation of the pre- to post-stimulation effects. **(a)** For the sham condition, the pre- to post-increase in alpha amplitude is a function of an increase in the aperiodic intercept alone. **(b)** The pre- to post-tACS effect reflects a combined increase of both the aperiodic intercept and oscillatory alpha activity.

Taken together, these results may suggest that the previously reported alpha power increase with time-on-task [39] is at least partly due to an increase in intercept and thus unrelated to oscillatory alpha activity. In the context of tACS aftereffects, this holds for both sham stimulation and tACS, with the latter producing an additional increase in oscillatory power. If the increase in intercept is relatively strong while the tACS effect is relatively weak, a true difference between sham stimulation and tACS might not be detected in 1/f-uncorrected data.

## 5 Conclusions

Spectral parametrization of the frequency spectrum into periodic and aperiodic components enables for a better characterization of tACS effects on oscillatory power. Conclusively, tACS seems to be frequency specific and does not affect the underlying aperiodic spectrum. However, this aperiodic component varies over time, signifying the importance of correcting for it when analyzing aftereffects of brain stimulation on brain oscillations.

## 6 Acknowledgements

This research was supported by the Neuroimaging Unit of the Carl von Ossietzky University Oldenburg funded by grants of from the German Research Foundation (3T MRI INST 184/152-1 FUGG and MEG INST 184/148-1 FUGG). Christoph S. Herrmann was supported by a grant of the German Research Foundation (Deutsche Forschungsgemeinschaft, DFG) under Germany’s Excellence Strategy – EXC 2177/1 - Project ID 390895286.

## 7 Author Contributions

FHK, RL, and CSH designed the study. FHK and RL carried out the analyses. All authors wrote the manuscript.

## 8 Conflict of interest

CSH holds a patent on brain stimulation. FHK, RL and DS, declare no competing interests.

